# Do Pseudosequences Matter in Neoantigen Prediction?

**DOI:** 10.64898/2025.12.09.693250

**Authors:** Adele Valeria, Rachel Karchin

## Abstract

Computational prediction of neoantigens that elicit T cell responses is central to the development of personalized cancer vaccines. Many current predictors represent MHC class I alleles using selected subsets of residues, known as pseudosequences, yet the extent to which pseudosequence choice and encoding strategy influence predictive performance has not been systematically examined. This study addresses that gap by evaluating a range of MHC representations within the BigMHC EL framework. We compared pseudosequence definitions based on protein structure and evolutionary diversity, a randomly sampled pseudosequence baseline, pseudosequences of varying lengths, embeddings generated using the ESM-2 protein language model, and a graph-based annotation embedding derived from allele groupings. Models using biologically informed pseudosequences consistently outperformed the random baseline, underscoring the importance of residue selection. Protein structure and evolutionary diversity pseudosequences showed similar performance, likely reflecting overlap in residues near the peptide-binding groove. We also found that pseudosequences of approximately 30 to 35 residues produced the strongest performance. Lastly, ESM-2 and annotation-based embeddings outperformed the random baseline but did not surpass curated pseudosequences under the current setup. Together, these findings indicate that curated pseudosequences remain efficient representations of MHC alleles in neoantigen prediction models, while alternative encodings can approximate but not yet replace residue-level sequence information.

## 1 Introduction

Personalized cancer immunotherapy has emerged as a pivotal approach in oncology, where treatments are tailored to the specific genetic alterations present in a patient’s tumor. This strategy builds on the central dogma of molecular biology, in which somatic mutations in tumor DNA are transcribed into RNA and translated into proteins that can be altered or entirely novel relative to those in healthy tissue. To prevent the accumulation of excess or dysfunctional proteins, cells undergo routine protein turnover, where intracellular proteins are degraded by the proteasome into short peptides. This process maintains cellular homeostasis and also drives immune surveillance through antigen presentation by major histocompatibility complex (MHC) molecules. Among the peptides generated through proteasomal degradation are those that originate from mutated tumor proteins, known as neoantigens (Schumacher and Schreiber 2015). These neoantigens can enter the MHC class I processing pathway and be presented on the tumor cell surface as peptide–MHC (pMHC) complexes, where they may be recognized by CD8^+^ T cells.

The recognition of tumor cells by T cells begins with the immune system continuously monitoring tissue to distinguish self from non-self for the detection of abnormal or transformed cells. When T cells encounter pMHC complexes, an immune response against cancer cells may be initiated if the presented peptide is recognized as foreign. Because healthy cells do not express these mutation-derived peptides, they are typically spared from immunemediated destruction. Neoantigens are therefore highly specific targets for personalized cancer immunotherapy. However, not all mutated peptides are immunogenic or capable of eliciting a functional T cell response. In practice, only a small fraction of mutated peptides are recognized by cytotoxic T cells. Clinical trials of personalized neoantigen vaccines in lung cancer (Ingels et al. 2024), melanoma (Ott et al. 2017), renal cell carcinoma (Braun et al. 2025), and pancreatic cancer (Sethna et al. 2025) have consistently shown that although many putative neoantigens can be computationally predicted, only a limited subset elicits detectable immune responses in vivo. This limited immunogenicity reflects multiple biological checkpoints a neoantigen must satisfy, including efficient intracellular processing (Abelin et al. 2017), stable presentation by MHC (Rasmussen et al. 2016), recognition by a pre-existing T cell receptor repertoire (Finnigan et al. 2024), and sufficient co-stimulatory and cytokine signaling within the tumor microenvironment (Shae et al. 2020; Li et al. 2021). Together, these constraints create a stringent bottleneck that narrows the pool of neoantigens capable of driving a productive antitumor immune response.

As a result, accurately identifying which mutated peptides are truly immunogenic remains a major challenge in cancer immunotherapy. Experimental identification of immunogenic neoantigens in individual patients typically relies on labor-intensive assays that are difficult to scale, limiting accessibility to neoantigen-based vaccines. To address this bottleneck, there has been growing interest in computational methods to predict and prioritize candidate neoantigens for downstream validation. Notable examples include BigMHC (Albert et al. 2023), DeepHLApan (Wu et al. 2019), NetMHCpan (Reynisson et al. 2020), MixMHCpred (Tadros et al. 2025), MHCflurry (O’Donnell et al. 2020), MHCnuggets (Shao et al. 2020), and PRIME (Gfeller et al. 2023). These models are trained on datasets that include both the amino acid sequence of the peptide and features of the corresponding MHC allele. Since MHC class I neoantigen peptides are short (typically 8 to 11 residues), their full sequences are used directly as input. In contrast, the MHC class I heavy chain sequence is substantially longer. For example, the HLA-A *α*-chain is 365 amino acids in length and includes the peptide-binding domains, a transmembrane segment, and a cytoplasmic tail. Thus, using the full MHC sequence as input may increase computational cost and introduce noise from regions not involved in peptide binding or presentation.

To address this limitation, many existing models use MHC pseudosequences, which are compact representations of selected residues believed to influence peptide binding and presentation. For example, BigMHC EL uses a 30-residue pseudosequence derived from the highest-entropy positions in a multiple sequence alignment of 18,929 MHC-I protein sequences from 12 species, assuming that the least conserved positions capture the variation most relevant for peptide binding and presentation. NetMHCpan-4.1 instead defines a 34-residue pseudosequence based on peptide-contacting positions identified from pMHC crystal structures, specifically residues within 4.0 Å of the bound peptide. Several prediction tools adopted variations of this representation. MHCflurry extends the NetMHCpan pseudosequence by adding three additional positions to improve discrimination among closely related alleles (O’Donnell et al. 2020), while PRIME retains the original 34 positions but encodes them with a BLOSUM62 substitution matrix (Gfeller et al. 2023). Other studies have instead explored using full MHC sequences. Recent transformer-based models such as PerceiverpMHC report performance comparable to pseudosequence-based models (Kushwaha et al. 2024), a finding that is consistent with at least one other full-sequence approach (Giziński et al. 2024). Together, these strategies highlight a lack of consensus regarding how MHC information should be represented and encoded.

These observations motivate a re-examination of the role of pseudosequences in neoantigen prediction, including how residue selection influences performance, how pseudosequence length should be considered, and when alternative representations may be advantageous. We examine three residue-selection strategies for defining pseudosequences, evaluate the influence of pseudosequence length, replace one-hot encodings with embeddings derived from ESM-2, and introduce an embedding based on nomenclature and standard HLA groupings. These groupings include P-groups, which cluster alleles with identical antigen-binding domain sequences, and HLA supertypes, which group alleles with similar peptide-binding properties based on shared binding motifs and similarity at key B- and F-pocket residues (Sidney et al. 2008), or on three-dimensional structural similarity of the peptide-binding groove (Shen et al. 2023). By training and benchmarking BigMHC EL models on each representation, we aim to characterize the contribution of residue selection and encoding to predictive performance and determine whether pseudosequences are necessary, or if simpler or alternative representations may be sufficient. In this study, the different MHC representations are compared within the BigMHC EL framework. We restricted our benchmarking to the BigMHC EL framework because it is fully open source and supports retraining with user-specified MHC representations. In contrast, NetMHCpan is a closed-source model that cannot be retrained or systematically modified to incorporate alternative pseudosequences. Similarly, while MHCflurry and PRIME are publicly available, their architectures and training pipelines are not readily configurable for repeated retraining across multiple pseudosequence variants as required in this study. Therefore, BigMHC EL provided the most practical and reproducible framework for controlled evaluation of MHC representations under identical training conditions.

## Methods

### Training and Evaluation Datasets

We used the same training, validation, and test datasets that were originally collected and partitioned for the development of BigMHC EL. These datasets are derived from publicly available single-allele immunopeptidomic experiments, in which cells are engineered to express a single HLA allele and incubated with a diverse peptide mixture. The resulting pMHC complexes are isolated and analyzed using liquid chromatography–mass spectrometry (LC– MS) to identify peptides presented on the cell surface (Abelin et al. 2017). This procedure produces eluted ligand (EL) datasets containing a ground truth set of positive pMHC pairs. Because LC–MS experiments do not directly identify non-presented peptides, negative examples were generated by sampling peptides from the human proteome that matched the expected length distribution of class I ligands.

The training dataset consisted of 288,032 positive and 16,739,285 negative pMHC pairs. These were split into approximately 90% for training and 10% for validation, yielding 259,298 positive and 15,065,287 negative examples in the training set (15,324,585 total) and 28,734 positive and 1,673,998 negative examples in the validation set (1,702,732 total). Both sets contained 149 HLA class I alleles. The held-out test set contained 45,409 positive and 900,592 negative pMHC pairs (946,001 total) spanning 36 alleles. All training and evaluation data is available at (https://data.mendeley.com/datasets/dvmz6pkzvb/4).

### MHC Pseudosequences

We represent each MHC class I allele using a pseudosequence, defined as a set of selected amino acid positions in the MHC sequence that are expected to influence peptide binding and presentation. We compared two established MHC class I pseudosequence definitions and included a random baseline. The BigMHC EL pseudosequences consist of 30 positions with the highest entropy in a multiple sequence alignment of MHC class I proteins across 12 species. The NetMHCpan-4.1 pseudosequences consist of 34 residues that directly contact the peptide in crystal structures, chosen as positions within approximately 4.0 ^°^A of the bound ligand. For the random baseline, we uniformly sampled 34 positions from all 364 alignment coordinates in the global multiple sequence alignment used to derive the BigMHC EL pseudosequences, thereby preserving the overall positional diversity of the default BigMHC EL residue set.

### Variable-Length BigMHC EL Pseudosequences

To assess how pseudosequence length affects model performance, we obtained the information content at each alignment position and ranked all 364 positions accordingly. For each *k* ∈ {5, 10, 15, 20, 25, 35, 50, 80, 100}, we selected the top *k* positions and extracted the corresponding residues for all alleles, yielding *k*-length pseudosequences that replaced the default 30-residue BigMHC EL pseudosequences.

### One-Hot Encoding

For each selected pseudosequence position, all amino acids observed at that position across alleles in the alignment were enumerated. Each amino acid was assigned a binary indicator column. Ambiguous residues (X) were treated as distinct categories. All binary indicators across positions were concatenated to form a fixed-length vector. For example, encoding the 30-residue BigMHC EL pseudosequence produced a 414-dimensional vector. These vectors were used as the MHC input to the neural network.

### ESM Embedding of MHC-I Pseudosequences

We embedded BigMHC EL and NetMHCpan-4.1 pseudosequences using the pretrained ESM-2 model (esm2 t33 650M UR50D). Pseudosequences were tokenized and processed in evaluation mode on a GPU. Hidden states from the final transformer layer were extracted, and embeddings corresponding to amino acid residues were mean-pooled to produce a 1280- dimensional vector per allele. These continuous vectors replaced the one-hot pseudosequences, and the BigMHC EL input layer was modified to accept float-valued inputs.

### Annotation-Based Embedding

We constructed an undirected graph in which each node corresponded to an HLA allele. Weighted edges were added between alleles that shared a supertype, P-group, two-field designation, or locus, with decreasing edge weights assigned in that order. Node2Vec (embedding dimension 64, walk length 12, 120 walks per node, *p* = 1, *q* = 1) was then used to generate continuous allele embeddings, which replaced pseudosequence encodings.

### Model Training and Selection

The original BigMHC EL ensemble was simplified by training models with three batch sizes (1,024, 4,096, and 16,384). A batch size of 4,096 was selected for all subsequent experiments. All models were trained for 50 epochs using the AdamW optimizer and binary cross-entropy loss with a learning rate of 5 × 10*^−^*^5^. Training was performed on the Rockfish computing cluster using NVIDIA A100 GPUs (2 GPUs, 24 CPUs, 96 GB RAM per run). A fixed random seed was used across experiments. After each epoch, validation AUPRC was computed, and the checkpoint with the highest validation AUPRC was selected for final testing.

## 2 Results

### 2.1 Biologically Informed Pseudosequences Outperform a Random Baseline

To evaluate how different pseudosequences affect neoantigen prediction, we trained BigMHC EL using protein-structure-based (as defined by NetMHCpan-4.1), evolutionary divergence-based (as defined by BigMHC EL), and randomly sampled pseudosequences, each encoded with one-hot representations (Methods). For each pseudosequence type, models were trained using three batch sizes (1,024, 4,096, and 16,384). Models trained with a batch size of 4,096 yielded AUROC and AUPRC values that were consistently comparable to, and in some cases slightly higher than, those of models trained at the other batch sizes (Figure not shown). Accordingly, the batch size of 4,096 was used for all subsequent benchmarking analyses.

When comparing MHC representations on the held-out test set, models using the BigMHC EL and NetMHCpan-4.1 pseudosequences achieved similar performance, with mean AUROC scores of 0.9675 and 0.9679 and mean AUPRC scores of 0.8245 and 0.8249 across batches, respectively. In contrast, the random baseline, generated by uniformly sampling 34 positions from the same multiple sequence alignment used for BigMHC EL, performed worse than the curated BigMHC EL and NetMHCpan-4.1 pseudosequences, with an AUROC of 0.9512 and an AUPRC of 0.7408. As shown in Figure 1A, the performance differences between the biologically informed pseudosequences and the random baseline were statistically significant (*p <* 0.001) at batch 4096, while the differences between the BigMHC EL and NetMHCpan-4.1 pseudosequences were negligible (Supplementary Information 1).

**Figure 1:**
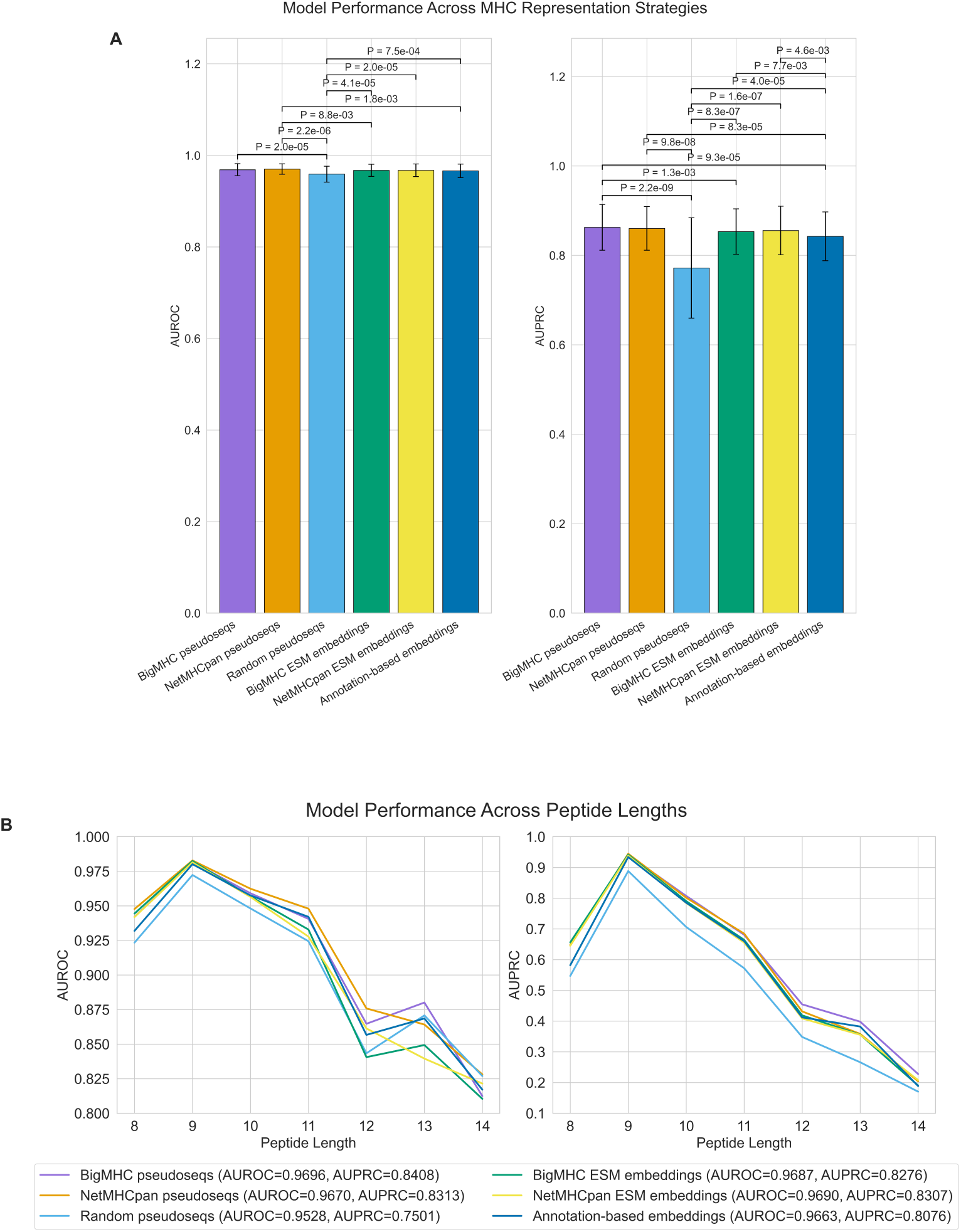
Comparison of model performance across different MHC representations. (A) Mean AUROC (left) and AUPRC (right) across alleles for models trained using six MHC representations. Error bars show standard deviation. Pairwise differences were tested using the Wilcoxon signed-rank test with Benjamini–Hochberg correction, and significant adjusted P-values (P ¡ 0.01) are shown above the bars. (B) Mean AUROC (left) and AUPRC (right) stratified by peptide length (8–14), averaged across alleles that contain both positive and negative peptides at each length.

To further assess model behavior, we stratified performance by peptide length (Figure 1B). Models with both NetMHCpan-4.1 and BigMHC EL pseudosequences maintained high AUROC and AUPRC values across peptides of length 8–14 and consistently exceeded the performance of the random baseline across all lengths. The highest performance for all models was observed for 9-mer peptides, which aligns with the well-established preference of MHC class I molecules for binding peptides of this length.

We next assessed each model’s ability to prioritize true presented peptides near the top of the ranked prediction list by computing precision among the top *n* predictions, i.e., positive predictive value (PPV) for the top *n*. This metric reflects the fraction of true positives within the highest-scoring *n* peptides. Models trained with both BigMHC EL and NetMHCpan-4.1 pseudosequences sustained precision values above 0.98 across the top 10,000 predictions, demonstrating strong ranking capabilities (Figure 2A). In contrast, the model trained on random pseudosequences showed lower and gradually declining precision with increasing *n*, indicating reduced ability to prioritize true presented peptides (Figure 2B).

**Figure 2:**
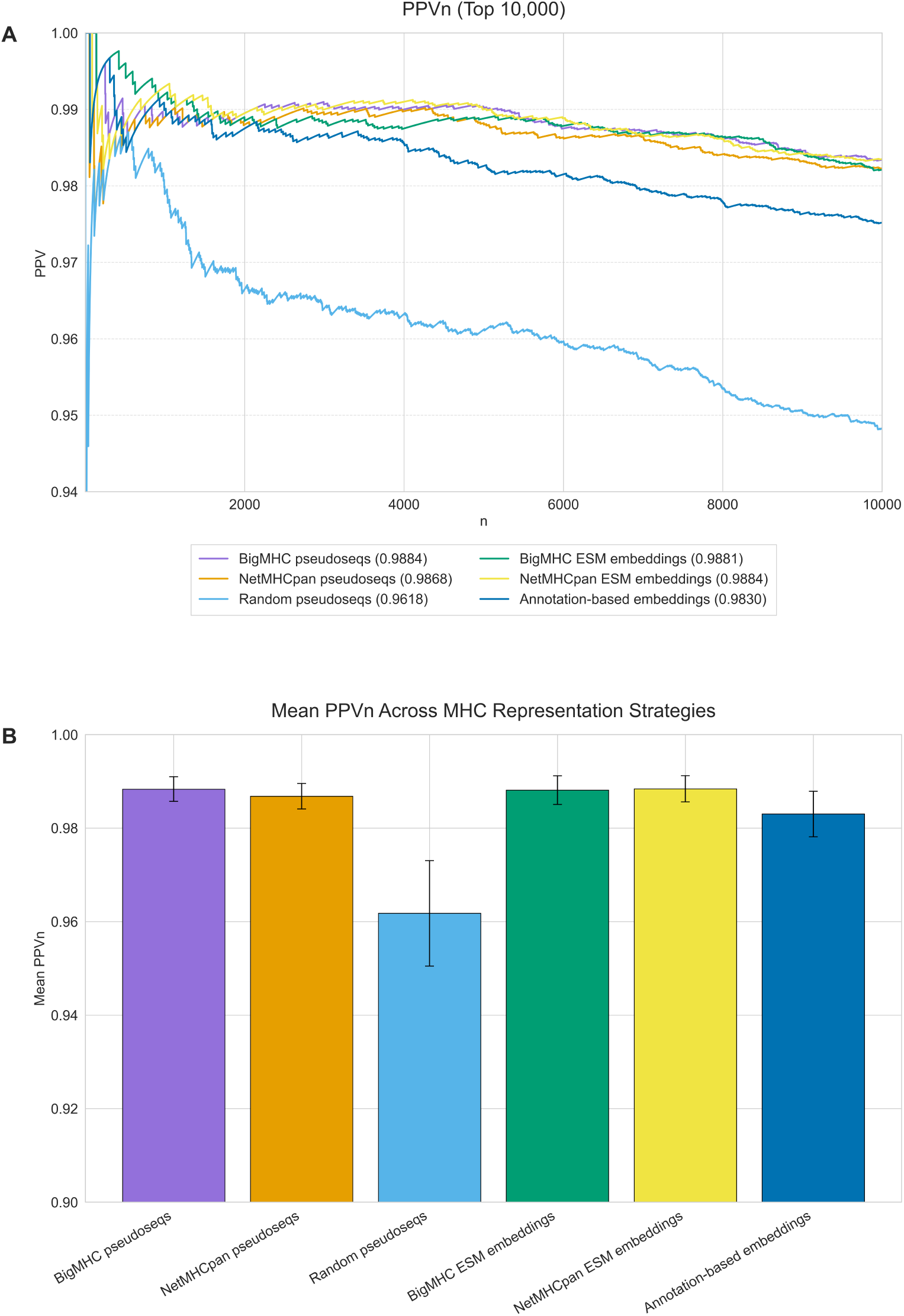
PPVn performance across MHC representations. (A) PPVn curves for each model, obtained by ranking all peptides by predicted score and computing PPV as a function of *n* over the top 10,000 predictions. The mean PPV across the top 10,000 ranks is shown in parentheses in the legend. (B) Mean PPVn across the top 10,000 ranked predictions for each model, with error bars representing the standard deviation of PPVn across ranks 1–10,000.

Similarly, per-locus and per-allele results showed consistent patterns across both AUROC and AUPRC. For the majority of HLA alleles, models using BigMHC EL and NetMHCpan-4.1 pseudosequences outperformed the random baseline (Figure 3A-B).

**Figure 3:**
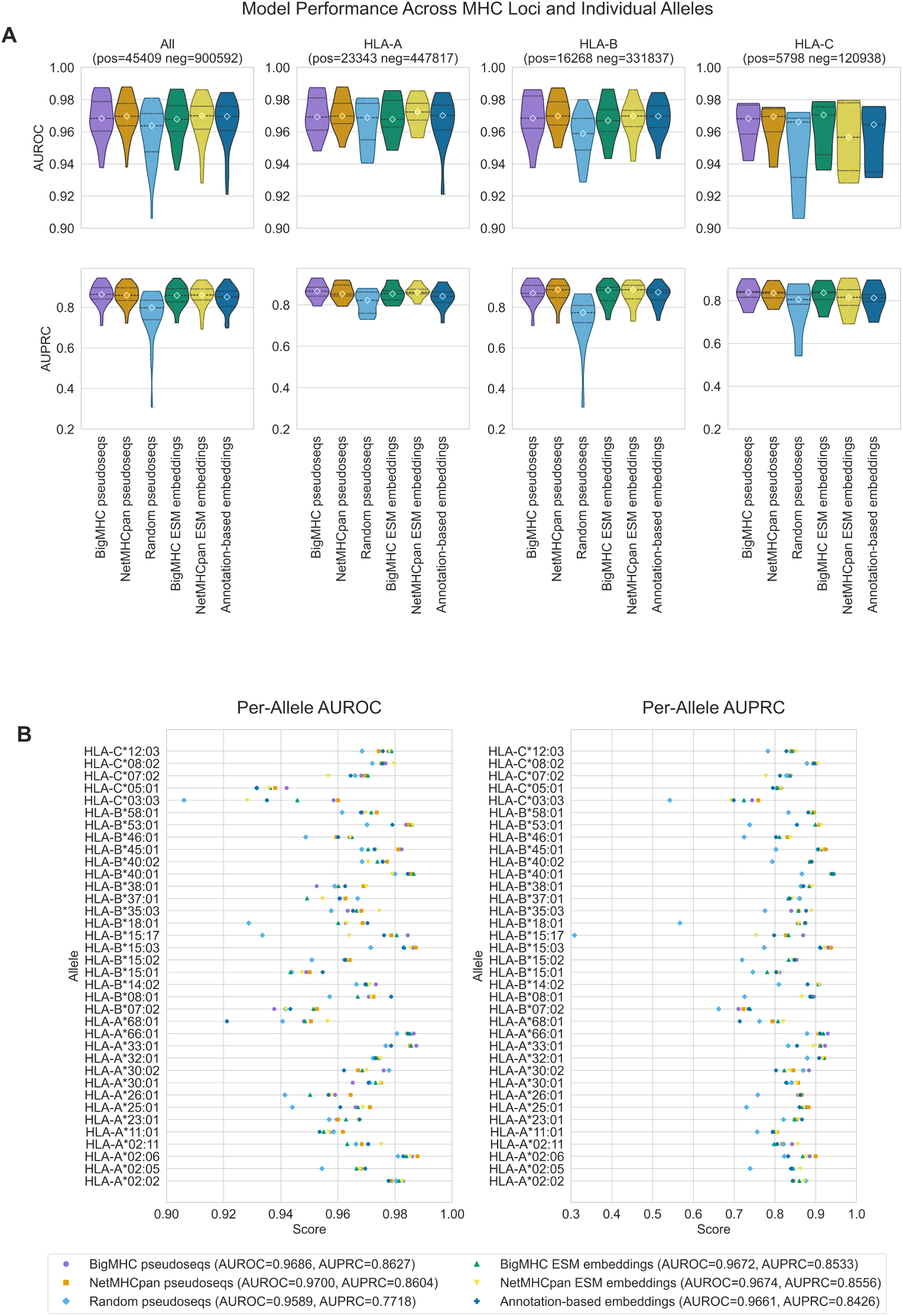
Model performance per-locus and per-allele across models. (A) Per-allele AUROC (top) and AUPRC (bottom) for six MHC representation models, shown across all alleles and separately for HLA-A, HLA-B, and HLA-C. Violin plots show the distribution across alleles, and diamonds mark the median for each model. (B) Per-allele AUROC and AUPRC displayed as scatter plots, where each point represents the performance of one model on one allele. Alleles lacking both positive and negative peptides were excluded.

**Figure 4:**
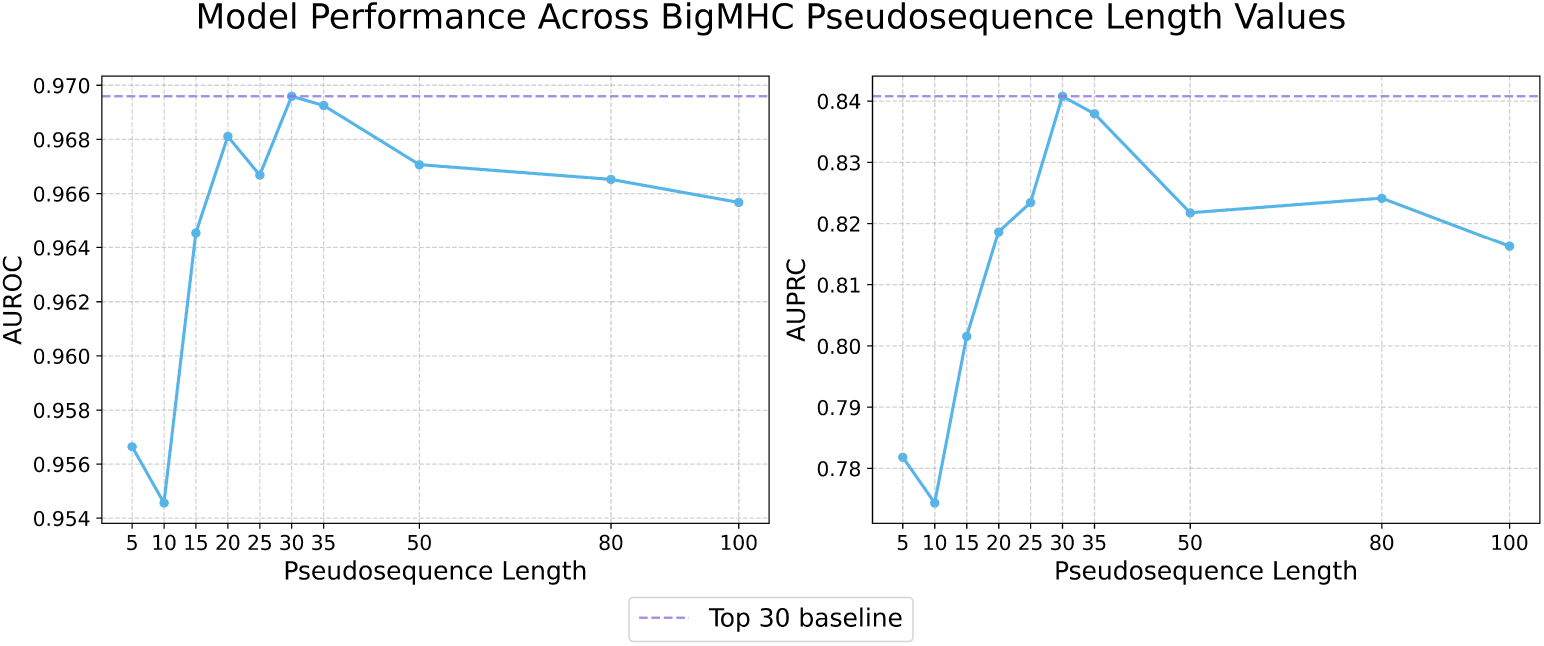
Model performance across BigMHC EL pseudosequence lengths. AU-ROC (left) and AUPRC (right) for models retrained using BigMHC EL pseudosequences of varying lengths. The dashed line marks performance at length 30, which corresponds to the original BigMHC EL pseudosequence and also yields the highest performance.

#### Effect of Pseudosequence Length on Prediction Performance

To evaluate how pseudosequence length influences predictive performance, we revisited the multiple sequence alignment used to construct the original BigMHC EL pseudosequence of 30 amino acid residues, calculated positional entropy for all alignment sites, and selected the top 5, 10, 15, 20, 25, 35, 50, 80, and 100 residues. Independent BigMHC EL models were then trained using each truncated or extended pseudosequence. Model accuracy generally increased with pseudosequence length up to 30 residues, after which performance declined (Figure 1A). The highest performance was observed at 30 residues, while the 35-residue model showed only a slight reduction in comparison. (Supplementary Information 2)

### 2.2 Annotation-Based Embedding

Our findings thus far indicate that pseudosequence-based representations effectively capture MHC sequence variation relevant to peptide binding and presentation. However, whether explicit sequence information is necessary for accurate prediction remains uncertain. To investigate this, we examined whether meaningful MHC representations could instead be derived from higher-level, annotation-based features. Specifically, we developed an annotation-based embedding that encodes relationships among MHC alleles according to shared P-group, HLA supertype, two-field nomenclature, and locus information.

We constructed a weighted graph in which nodes represent alleles and edges connect alleles sharing any of these attributes. When two alleles shared multiple attributes, their edge weights were summed accordingly. The two-field and locus attributes also served as fallback connections to ensure coverage of alleles lacking P-group or supertype annotations. Using this graph, we applied node2vec to learn embeddings of MHC alleles based on their shared annotation-derived relationships (Grover and Leskovec 2016).

The annotation-based embedding model achieved an AUROC of 0.9663 and an AUPRC of 0.8076 on the held-out test set, outperforming the random pseudosequence model (Figure 1A). These results suggest that residue-level sequence variation likely contributes additional information that is not fully captured by higher-level annotations. Nonetheless, the relatively strong performance of the annotation-based model indicates that part of this information can be approximated through structured features, suggesting that explicit sequence input may not always be strictly required for achieving competitive predictive performance.

#### 2.2.1 ESM-Based Embedding

For each pseudosequence type, residue-level embeddings were extracted using the ESM-2 model (650M parameters) and mean-pooled across sequence positions to obtain a fixed-length vector of 1,280 dimensions per allele. These embeddings were then used to replace the one-hot encoded pseudosequences as the MHC input. Both BigMHC EL and NetMHCpan-4.1 pseudosequences encoded with ESM embeddings achieved highly similar performance, with the BigMHC–ESM model reaching an AUROC of 0.9687 and AUPRC of 0.8276, and the NetMHCpan–ESM model achieving an AUROC of 0.9690 and AUPRC of 0.8307. The differences between ESM-based and one-hot encoded models were not statistically significant.

## 3 Discussion

In this study, we evaluated how different representations of MHC class I molecules influence prediction performance within the BigMHC EL framework. Although neoantigen predictors commonly rely on compact residue subsets to encode MHC polymorphism, the extent to which residue choice, pseudosequence length, and encoding strategy affect model accuracy has remained unclear. Our results point to three main observations. First, curated pseudosequences derived from structural or evolutionary criteria remained the most reliable representations under matched training conditions, suggesting that biologically informed residue selection remains important. Second, a compact window of roughly 30 to 35 residues appeared to capture much of the informative variation required for accurate prediction. Third, alternative encodings such as ESM-based embeddings and an annotation-derived graph embedding captured some aspects of MHC variation but did not match the accuracy achieved by curated pseudosequences in our setup.

In the first set of experiments, where we compared curated pseudosequences with a randomly sampled baseline, we observed that the BigMHC EL pseudosequences- and NetMHCpan-4.1 pseudosequences-based models achieved similar performance despite being defined using different selection principles. A likely explanation is their substantial overlap in the residues chosen for each pseudosequence. When mapped onto the multiple sequence alignment used to derive the BigMHC EL pseudosequences, the two strategies share 18 positions clustered within the *α*_1_ and *α*_2_ domains that form the peptide-binding groove. These shared sites likely contribute a significant portion of the predictive signal.

This convergence also suggests that whether residues are selected based on structural proximity to the ligand or on sequence entropy across alleles and species, both strategies ultimately emphasize many of the same biologically important positions. In other words, the least conserved positions of MHC class I across alleles and species may be precisely those that shape peptide-binding preferences, making them more likely to be captured through structural criteria accordingly. Another potential explanation for the similar performance is the relatively small difference in pseudosequence length between the BigMHC EL (30 residues) and NetMHCpan-4.1 (34 residues) definitions. In the subsequent length-sweep experiments, where we varied the number of residues included in the BigMHC EL pseudosequences, model performance increased as key polymorphic positions were added, reached a maximum at 30 residues, and then plateaued before declining once positions beyond this range introduced more noise than informative signal. Given that both BigMHC EL and NetMHCpan-4.1 pseudosequences fall within this approximate optimal window of 30 to 35 residues, their similarly strong predictive performance aligns with the length-dependent trends observed.

We also examined whether replacing one-hot encodings of pseudosequences with embeddings derived from large protein language models would provide any benefit. Using mean-pooled, frozen ESM-2 representations resulted in performance comparable to models trained with BigMHC EL and NetMHCpan-4.1 one-hot pseudosequences. The reason the embeddings did not improve performance is not entirely clear. One possibility is that one-hot inputs already isolate the key polymorphic positions, which may limit the ability of learned or pretrained embeddings to introduce additional signal.

Another consideration is methodological. Our analysis focused specifically on encoding strategies under matched downstream architectures, so ESM-2 was used only as a frozen feature extractor with standard pooling. We did not apply fine-tuning or explore alternative pooling strategies, such as structural- or pocket-aware pooling that can assign greater weight to residues forming the peptide-binding groove. However, these approaches would introduce model changes that extend beyond a direct comparison of pseudosequence encodings and are more appropriate for future work aimed at developing augmented models.

Finally, we asked whether pseudosequences are strictly necessary. To examine this, we moved beyond direct MHC sequence representations and constructed an annotation-based embedding that captures higher-level, biologically informed groupings such as shared P-group, HLA supertype, and allele nomenclature. These groupings are ultimately based on sequence or structural similarity, yet they avoid using explicit residue identities. This analysis showed that competitive performance does not strictly require residue-level input. The annotation graph captured enough allele relationships to outperform a random baseline, although it did not reach the performance of curated pseudosequences.

Most of the graph structure came from nomenclature attributes rather than supertypes or P-groups. Among the 149 alleles in our datasets, only 299 edges were contributed by P-group or supertype membership. The remaining alleles were connected through two-field or locus attributes, which contributed 3,321 fallback edges and ensured full connectivity. These results suggest that allele identity, as encoded by the allele name and its hierarchical structure, carries meaningful biological information. Nonetheless, current binding predictors make limited use of allele names as explicit features, and to the best of our knowledge none has isolated the contribution of allele labels independently of sequence- or pseudosequence-based representations.

Our current annotation-based embedding serves as an initial exploration of how far allele names and their associated groupings can be leveraged. Future work could extend this approach by incorporating additional biological relationships, experimenting with alternative graph construction strategies, or evaluating whether annotation-based embeddings can complement pseudosequences to further improve BigMHC EL performance. At the same time, the remaining performance gap suggests that residue-level variation captures information that higher-level annotations alone cannot yet fully recover.

All of these findings, however, should be interpreted with caution. Our training setup used a simplified version of the original BigMHC EL ensemble, and larger-scale training may reveal differences not captured here. Moreover, all experiments were conducted within the BigMHC EL architecture and training protocol, leaving open the question of whether the same trends would extend to other model families. The pseudosequence length analysis also relied specifically on the BigMHC EL entropy-based residue set, and it remains uncertain whether similar patterns would emerge under alternative residue-selection strategies. Finally, the training data were derived from eluted-ligand experiments with sampled negatives, and we did not conduct leave-allele-out or prospective evaluations on alleles absent from training. In conclusion, these observations have practical implications for selecting an MHC allele representation. When allele sequences are available and performance is the primary objective, a curated 30–35-residue pseudosequence with position-specific encoding remains the most reliable option. It consistently yielded the strongest results in our evaluations and is straightforward to implement, making it suitable for both training and deployment. When allele sequences are unavailable, incomplete, or inconsistent, a structured annotation embedding can serve as an alternative. Although less accurate than sequence-based representations, it nevertheless captured sufficient relationships to produce coherent rankings. Protein language model features represent another potential approach, particularly when a richer sequence representation is desired. In our experiments, however, frozen, mean-pooled PLM embeddings did not improve performance relative to one-hot pseudosequences. Any meaningful benefit will likely require fine-tuning or pooling strategies that emphasize functionally relevant regions rather than relying on global averages.

## Availability

All code and data used in this study including Supplementary Information are available on Github (https://github.com/KarchinLab/Do-Pseudoseqs-Matter) and Mendeley Data (https://doi.org/10.17632/pn8zym54pz.1).

## Author Contributions

A.V. and R.K. conceived the project and designed the study. A.V. developed the computational pipeline. A.V. and R.K. validated the analysis and verified its reproducibility. A.V. wrote the first draft of the manuscript. A.V. and R.K. reviewed and edited the manuscript. R.K. supervised the project.

## Acknowledgments

We thank Benjamin Albert for valuable discussions. This work was supported by the National Institutes of Health [grant number R01CA301054 to R.K.].

## References

Abelin, J. G., Keskin, D. B., Sarkizova, S., Hartigan, C. R., Zhang, W., Sidney, J., Stevens, J., Lane, W., Zhang, G. L., Eisenhaure, T. M., Clauser, K. R., Hacohen, N., Rooney, M. S., Carr, S. A., and Wu, C. J. (2017). Mass spectrometry profiling of HLA-associated peptidomes in mono-allelic cells enables more accurate epitope prediction. 46(2):315–326.

Albert, B. A., Yang, Y., Shao, X. M., Singh, D., Smith, K. N., Anagnostou, V., and Karchin, R. (2023). Deep neural networks predict class i major histocompatibility complex epitope presentation and transfer learn neoepitope immunogenicity. 5(8):861–872.

Braun, D. A., Moranzoni, G., Chea, V., McGregor, B. A., Blass, E., Tu, C. R., Vanasse, A. P., Forman, C., Forman, J., Afeyan, A. B., Schindler, N. R., Liu, Y., Li, S., Southard, J., Chang, S. L., Hirsch, M. S., LeBoeuf, N. R., Olive, O., Mehndiratta, A., Greenslade, H., Shetty, K., Klaeger, S., Sarkizova, S., Pedersen, C. B., Mossanen, M., Carulli, I., Tarren, A., Duke-Cohan, J., Howard, A. A., Iorgulescu, J. B., Shim, B., Simon, J. M., Signoretti, S., Aster, J. C., Elagina, L., Carr, S. A., Leshchiner, I., Getz, G., Gabriel, S., Hacohen, N., Olsen, L. R., Oliveira, G., Neuberg, D. S., Livak, K. J., Shukla, S. A., Fritsch, E. F., Wu, C. J., Keskin, D. B., Ott, P. A., and Choueiri, T. K. (2025). A neoantigen vaccine generates antitumour immunity in renal cell carcinoma. 639(8054):474–482.

Finnigan, J. P., Newman, J. H., Patskovsky, Y., Patskovska, L., Ishizuka, A. S., Lynn, G. M., Seder, R. A., Krogsgaard, M., and Bhardwaj, N. (2024). Structural basis for self-discrimination by neoantigen-specific TCRs. 15(1):2140.

Gfeller, D., Schmidt, J., Croce, G., Guillaume, P., Bobisse, S., Genolet, R., Queiroz, L., Cesbron, J., Racle, J., and Harari, A. (2023). Improved predictions of antigen presentation and TCR recognition with MixMHCpred2.2 and PRIME2.0 reveal potent SARS-CoV-2 CD8+ t-cell epitopes. 14(1):72–83.e5.

Giziński, S., Preibisch, G., Kucharski, P., Tyrolski, M., Rembalski, M., Grzegorczyk, P., and Gambin, A. (2024). Enhancing antigenic peptide discovery: Improved MHC-i binding prediction and methodology. 224:1–9.

Grover, A. and Leskovec, J. (2016). node2vec: Scalable feature learning for networks. In Proceedings of the 22nd ACM SIGKDD International Conference on Knowledge Discovery and Data Mining, pages 855–864. ACM.

Ingels, J., De Cock, L., Stevens, D., Mayer, R. L., Théry, F., Sanchez, G. S., Vermijlen, D., Weening, K., De Smet, S., Lootens, N., Brusseel, M., Verstraete, T., Buyle, J., Van Houtte, E., Devreker, P., Heyns, K., De Munter, S., Van Lint, S., Goetgeluk, G., Bonte, S., Billiet, L., Pille, M., Jansen, H., Pascal, E., Deseins, L., Vantomme, L., Verdonckt, M., Roelandt, R., Eekhout, T., Vandamme, N., Leclercq, G., Taghon, T., Kerre, T., Vanommeslaeghe, F., Dhondt, A., Ferdinande, L., Van Dorpe, J., Desender, L., De Ryck, F., Vermassen, F., Surmont, V., Impens, F., Menten, B., Vermaelen, K., and Vandekerckhove, B. (2024). Neoantigen-targeted dendritic cell vaccination in lung cancer patients induces long-lived t cells exhibiting the full differentiation spectrum. 5(5):101516.

Kushwaha, A., Duroux, P., Giudicelli, V., Todorov, K., and Kossida, S. (2024). IMGT/RobustpMHC: robust training for class-i MHC peptide binding prediction. 25(6):bbae552.

Li, Y., Cong, Y., Jia, M., He, Q., Zhong, H., Zhao, Y., Li, H., Yan, M., You, J., Liu, J., Chen, L., Hang, H., and Wang, S. (2021). Targeting IL-21 to tumor-reactive t cells enhances memory t cell responses and anti-PD-1 antibody therapy. 12(1):951.

Ott, P. A., Hu, Z., Keskin, D. B., Shukla, S. A., Sun, J., Bozym, D. J., Zhang, W., Luoma, A., Giobbie-Hurder, A., Peter, L., Chen, C., Olive, O., Carter, T. A., Li, S., Lieb, D. J., Eisenhaure, T., Gjini, E., Stevens, J., Lane, W. J., Javeri, I., Nellaiappan, K., Salazar, A. M., Daley, H., Seaman, M., Buchbinder, E. I., Yoon, C. H., Harden, M., Lennon, N., Gabriel, S., Rodig, S. J., Barouch, D. H., Aster, J. C., Getz, G., Wucherpfennig, K., Neuberg, D., Ritz, J., Lander, E. S., Fritsch, E. F., Hacohen, N., and Wu, C. J. (2017). An immunogenic personal neoantigen vaccine for patients with melanoma. 547(7662):217–221.

O’Donnell, T. J., Rubinsteyn, A., and Laserson, U. (2020). MHCflurry 2.0: Improved pan-allele prediction of MHC class i-presented peptides by incorporating antigen processing. 11(1):42–48.e7.

Rasmussen, M., Fenoy, E., Harndahl, M., Kristensen, A. B., Nielsen, I. K., Nielsen, M., and Buus, S. (2016). Pan-specific prediction of peptide–MHC class i complex stability, a correlate of t cell immunogenicity. 197(4):1517–1524.

Reynisson, B., Alvarez, B., Paul, S., Peters, B., and Nielsen, M. (2020). NetMHCpan-4.1 and NetMHCIIpan-4.0: improved predictions of MHC antigen presentation by concurrent motif deconvolution and integration of MS MHC eluted ligand data. 48:W449–W454.

Schumacher, T. N. and Schreiber, R. D. (2015). Neoantigens in cancer immunotherapy. 348(6230):69–74.

Sethna, Z., Guasp, P., Reiche, C., Milighetti, M., Ceglia, N., Patterson, E., Lihm, J., Payne, G., Lyudovyk, O., Rojas, L. A., Pang, N., Ohmoto, A., Amisaki, M., Zebboudj, A., Odgerel, Z., Bruno, E. M., Zhang, S. L., Cheng, C., Elhanati, Y., Derhovanessian, E., Manning, L., Müller, F., Rhee, I., Yadav, M., Merghoub, T., Wolchok, J. D., Basturk, O., Gönen, M., Epstein, A. S., Momtaz, P., Park, W., Sugarman, R., Varghese, A. M., Won, E., Desai, A., Wei, A. C., D’Angelica, M. I., Kingham, T. P., Soares, K. C., Jarnagin, W. R., Drebin, J., O’Reilly, E. M., Mellman, I., Sahin, U., Türeci, , Greenbaum, B. D., and Balachandran, V. P. (2025). RNA neoantigen vaccines prime long-lived CD8+ t cells in pancreatic cancer. 639(8056):1042–1051.

Shae, D., Baljon, J. J., Wehbe, M., Christov, P. P., Becker, K. W., Kumar, A., Suryadevara, N., Carson, C. S., Palmer, C. R., Knight, F. C., Joyce, S., and Wilson, J. T. (2020). Co-delivery of peptide neoantigens and stimulator of interferon genes agonists enhances response to cancer vaccines. 14(8):9904–9916.

Shao, X. M., Bhattacharya, R., Huang, J., Sivakumar, I. A., Tokheim, C., Zheng, L., Hirsch, D., Kaminow, B., Omdahl, A., Bonsack, M., Riemer, A. B., Velculescu, V. E., Anagnostou, V., Pagel, K. A., and Karchin, R. (2020). High-throughput prediction of MHC class i and II neoantigens with MHCnuggets. 8(3):396–408.

Shen, Y., Parks, J. M., and Smith, J. C. (2023). HLA class i supertype classification based on structural similarity. 210(1):103–114.

Sidney, J., Peters, B., Frahm, N., Brander, C., and Sette, A. (2008). HLA class i supertypes: a revised and updated classification. 9(1):1.

Tadros, D. M., Racle, J., and Gfeller, D. (2025). Predicting MHC-i ligands across alleles and species: how far can we go? 17(1):25.

Wu, J., Wang, W., Zhang, J., Zhou, B., Zhao, W., Su, Z., Gu, X., Wu, J., Zhou, Z., and Chen, S. (2019). DeepHLApan: A deep learning approach for neoantigen prediction considering both HLA-peptide binding and immunogenicity. 10:2559.

